# Actin turnover protects the cytokinetic contractile ring from structural instability

**DOI:** 10.1101/2022.02.23.481727

**Authors:** Zachary McDargh, Tianyi Zhu, Hongkang Zhu, Ben O’Shaughnessy

**Affiliations:** Department of Chemical Engineering, Columbia University, New York, NY 10027, USA

**Keywords:** cytokinesis, contractile ring, ADF/cofilin, actin turnover, instability

## Abstract

In common with other actomyosin contractile cellular machineries, actin turnover is required for normal function of the cytokinetic contractile ring. Cofilin is an actin-binding protein contributing to turnover by severing actin filaments, required for cytokinesis by many organisms. In fission yeast cofilin mutants, contractile rings suffer bridging instabilities in which actin bundles peel away from the plasma membrane into straight bridges. The origin of this behaviour is unclear. Here we used molecularly explicit simulations of the fission yeast contractile ring to examine the role of cofilin. Simulations reproduced the experimentally observed cycles of bridging and reassembly during constriction, each lasting ∼ 6 min, and the tendency for bridging to occur in ring segments with low myosin II Myo2 density. The lack of cofilin severing produced ∼ 2-fold longer filaments and, consequently, ∼ 2-fold higher ring tensions. Simulations identified bridging as originating in the boosted ring tension, which increased centripetal forces that detached actin from Myo2 that anchored actin to the membrane. Thus, cofilin serves a critical role in cytokinesis by protecting the contractile ring from bridging, the principal structural threat.

**Summary statement:** Molecularly explicit simulations showed that cofilin-mediated actin severing protects the fission yeast cytokinetic contractile ring from instabilities in which actin peels away into straight bridges.

## Introduction

Turnover of actin and other components is a universal feature of actomyosin contractile machineries in cells. Actin turns over through nucleation of filamentous actin by formins, Arp2/3 or other factors (Pollard, 2007), and by dissociation and/or disassembly of filaments. Cofilin is an actin-binding protein whose severing activity is critical for actin turnover in many machineries (Elam et al., 2013), including the actomyosin contractile ring that drives cell division during cytokinesis (D’Avino et al., 2015; Pollard and O’Shaughnessy, 2019; Pollard and Wu, 2010a). Cofilin is required for cytokinesis in many organisms (Abe et al., 1996; Gunsalus et al., 1995; Ono et al., 2003) including fission yeast (Nakano and Mabuchi, 2006), but many aspects of the mechanisms that make cofilin essential to cytokinesis remain poorly understood.

Fission yeast assembles a contractile ring from several hundred precursor protein complexes called nodes, each node incorporating myosin II Myo2, formin Cdc12 that nucleates and grows actin filaments, and other components (Wu et al., 2006). Normal ring assembly requires cofilin, as shown by studies of cytokinesis with mutants (Chen and Pollard, 2011; Nakano and Mabuchi, 2006). In cofilin mutants with reduced actin binding and severing activity, contractile rings failed to assemble normally from nodes, but instead assembled rings more slowly via diverse intermediate structures (Chen and Pollard, 2011). It was argued (Chen and Pollard, 2011) that cofilin-mediated actin filament severing is required for assembly from the nodes which are pulled together by pairwise actin filament connections that need regular renewal by severing to avoid node clumping and assembly failure (Vavylonis et al., 2008).

The role of cofilin during constriction of the assembled ring is much less clear. Is cofilin needed for constriction, and if so what is the mechanism? Fission yeast contractile rings with cofilin mutants constricted at normal mean rates, but with more variability (Chen and Pollard, 2011). However, during constriction rings exhibited remarkable structural irregularities, in which segments of the ring peeled away from the plasma membrane into straight segments we call bridges (Cheffings et al., 2019; Chen and Pollard, 2011). As bridging tended to occur in ring segments with lower Myo2 density, it was suggested the instability originated in a combination of reduced local Myo2-mediated actin anchoring and reduced local ring tension in the bridging region (Cheffings et al., 2019).

The contractile ring is a complex cellular machine of many components, so understanding the mechanisms of constriction and force production can benefit from mathematical modeling (Glotzer, 2005; O’Shaughnessy and Thiyagarajan, 2018; Pollard and O’Shaughnessy, 2019). Fission yeast is uniquely positioned for realistic quantitative modeling due to a wealth of available data on the identities, amounts, biochemical properties and organization of ring components (Chatterjee and Pollard, 2019; Chen et al., 2015; Courtemanche et al., 2016; Friend et al., 2018; Hayakawa et al., 2020; Pollard et al., 2017; Pollard and Wu, 2010a; Stark et al., 2010; Takaine et al., 2015; Wu and Pollard, 2005). This data was used to constrain models of the contractile ring which reproduced structural features and ring tensions measured in live cells (Alonso-Matilla et al., 2019; McDargh et al., 2021; Stachowiak et al., 2014; Thiyagarajan et al., 2017). In contrast to the classic sarcomeric force production mechanism of striated muscle (Huxley and Niedergerke, 1954; Huxley and Hanson, 1954), a sliding node mechanism emerged, where Myo2 and the unconventional myosin Myp2 (Bezanilla et al., 1997) pull nodes around the ring via attached actin filaments barbed-end-anchored to nodes via formins (McDargh et al., 2021). This generates tension in filaments, and a net ring tension. Experimental and modeling studies of septation, the growth of new cell wall in the wake of the constricting ring, suggest septation determines the constriction rate while the centripetal force from the ring mechanically regulates septation (Thiyagarajan et al., 2015; Zhou et al., 2015).

Here we used mathematical modeling to study the role of cofilin-mediated turnover in constricting contractile rings. Using a well-tested simulation framework (McDargh et al., 2021), we studied fission yeast contractile rings with the ADF-M3 cofilin mutant with reduced actin binding and severing activity (Chen and Pollard, 2011). Simulations quantitatively reproduced the experimentally observed bridges, the correlation with lower Myo2 densities and, remarkably, the cyclic character of bridging with ∼ 6 min per bridging-rehealing episode (Cheffings et al., 2019). The origin of the bridging instability is that actin filaments grow much longer without cofilin-mediated severing, so the ring tension is massively boosted, sufficient to pull actin filaments away from the Myo2 that anchors them to the membrane. In live cells (Laplante et al., 2015) and in simulations (McDargh et al., 2021), essentially the same bridging instability is seen in *myo2-E1* mutants with weakened binding to actin, showing that either increased centripetal loading or reduced anchoring provokes bridging. Thus, by maintaining actin filament lengths within a safe bound, cofilin protects the contractile ring from bridging instability, the principal structural threat to the contractile ring.

## Results

### Molecularly detailed simulation of the fission yeast cytokinetic ring

To examine the role of cofilin we adapted a molecularly explicit model of the fission yeast contractile ring we developed previously (McDargh et al., 2021), severely constrained by extensive data making the fission yeast contractile ring presently the most amenable to realistic mathematical modeling. The amounts of many ring components were measured over time and many were biochemically characterized (Chatterjee and Pollard, 2019; Chen et al., 2015; Courtemanche et al., 2016; Friend et al., 2018; Hayakawa et al., 2020; Pollard et al., 2017; Pollard and Wu, 2010b; Stark et al., 2010; Takaine et al., 2015; Wu and Pollard, 2005). Super-resolution FPALM microscopy showed the actomyosin organization in constricting rings is built from protein complexes called nodes (Laplante et al., 2016), anchored to the plasma membrane (PM). Each node contains myosin II Myo2, the actin filament nucleator formin Cdc12 and other components. A second unconventional myosin II Myp2 does not belong to the nodes and is likely unanchored from the plasma membrane.

Here we summarize the main points of the model (for more details see ref. (McDargh et al., 2021). Parameter values are listed in Material and Methods. In the model each node is a coarse-grained representation of the organization revealed by FPALM, with Myo2, formin Cdc12, IQGAP Rng2 and F-BAR protein Cdc15 (Laplante et al., 2016) (Fig. 1A). Formin Cdc12 dimers nucleate and grow actin filaments, binding the node 44 nm from the membrane, while the heads of ∼ 8 Myo2 dimers are represented by an ellipsoid with dimensions and location matching experiment (132 × 102 × 102 nm, 94 nm from the membrane). Semiflexible actin filaments are represented by chains of rods with bending stiffness corresponding to the ∼ 10 μm persistence length (Ott et al., 1993), and Myp2 is represented as unanchored 200 nm clusters of 16 molecules each (Cheffings et al., 2019; Laplante et al., 2015; Takaine et al., 2015).

**Figure 1.**
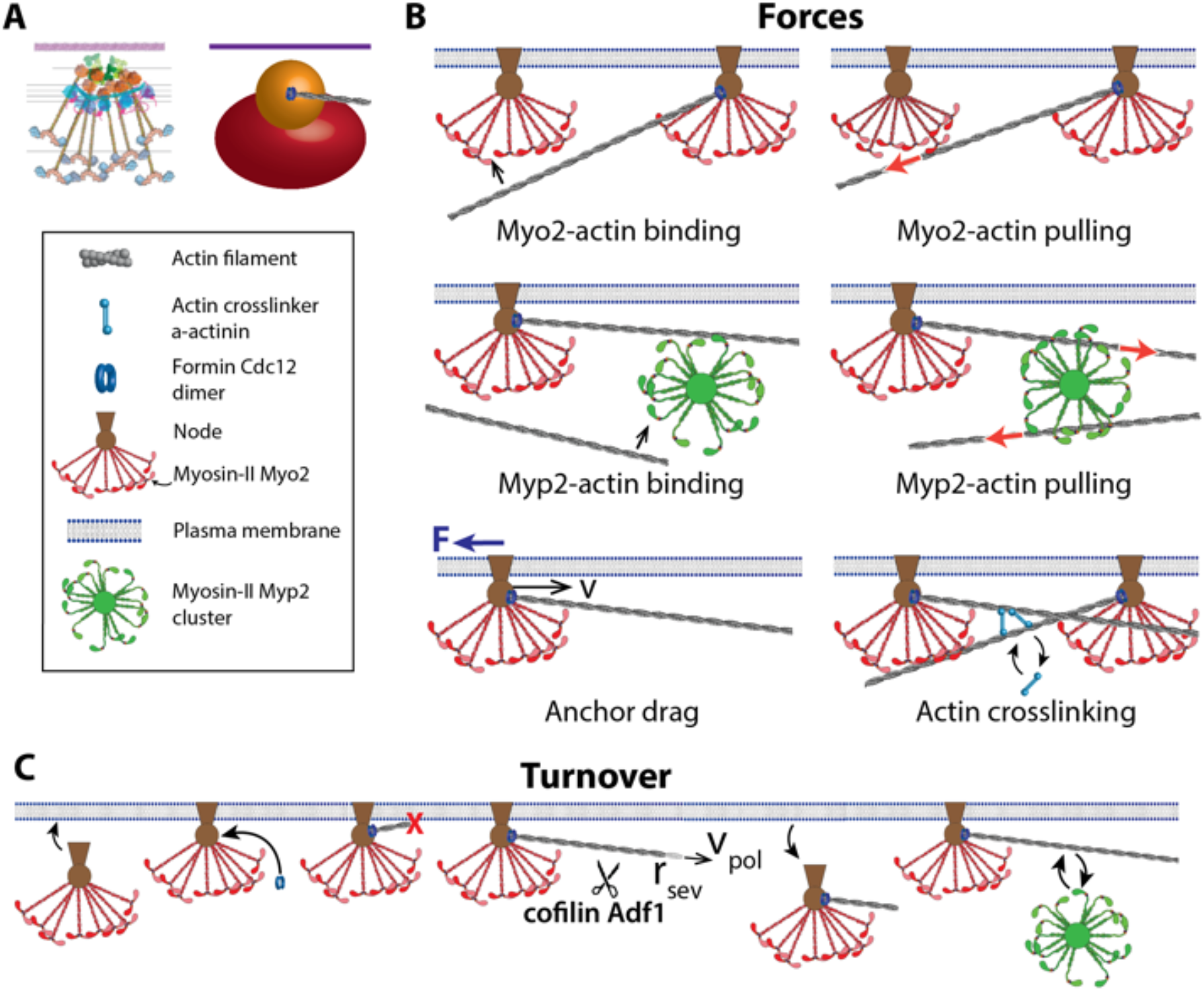
Molecularly explicit model of the *S. pombe* contractile ring. (A) Left: schematic of membrane-anchored node organization. Right: coarse-grained representation of a node used in the model. Myosin II Myo2 (red), formin Cdc15 (blue), IQGAP Rng2 (orange). (B) Forces in the model. (C) Turnover of ring components. Nodes bind and unbind the plasma membrane. Formin dimers bind nodes and polymerize randomly oriented actin filaments at rate *ν*_pol_. Cofilin Adf1 (scissors) stochastically severs filaments at rate *r*_sev_. Myp2 clusters bind and unbind actin filaments.

Filaments intersecting Myo2 or Myp2 clusters bind and are pulled (Fig. 1B), following linear force-velocity relations with total stall forces per cluster, 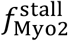 and 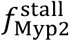, and with the measured load-free velocity 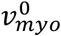 for the estimated 25 myosin II heads per actin filament, 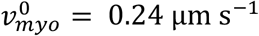 (Stark et al., 2010). Drag forces resist node membrane anchor motion with drag coefficient chosen to reproduce the experimental node velocity, 22 ± 10 nm s^−1^ (Laplante et al., 2016). Steric forces oppose overlap of components, ensuring actin filaments do not cross. Actin filaments are dynamically crosslinked by Ain1 α-actinin dimers (Wu and Pollard, 2005).

The component amounts follow the experimental values throughout constriction (Courtemanche et al., 2016; Wu and Pollard, 2005) (e.g., 3300 Myo2 molecules, 230 formin dimers in ∼ 210 nodes, 2300 Myp2 molecules in ∼ 140 clusters, and 230 actin filaments of total length ∼ 580 μm at constriction onset). Nodes stochastically bind a 0.2 μm wide zone of the ingrowing septum (Cortes et al., 2007), with mean dissociation time 41 s consistent with reported component dissociation times (Clifford et al., 2008; Pelham and Chang, 2002; Yonetani et al., 2008).

On binding a node a formin nucleates a randomly oriented actin filament. Cofilin-mediated severing stochastically shortens filaments homogenously with rate *r*_sev_ = 0.93 μm^−1^ min^−1^ (see Table 1) (Elam et al., 2013). Myp2 clusters bind actin filaments with turnover time 46 s (Takaine et al., 2015). Rates of severing, binding and actin polymerization are set by demanding simulations reproduce the experimental densities of Myo2, Myp2 and formin and mean actin length, all evolving as constriction progresses (Fig. 1C).

**Table 1.**
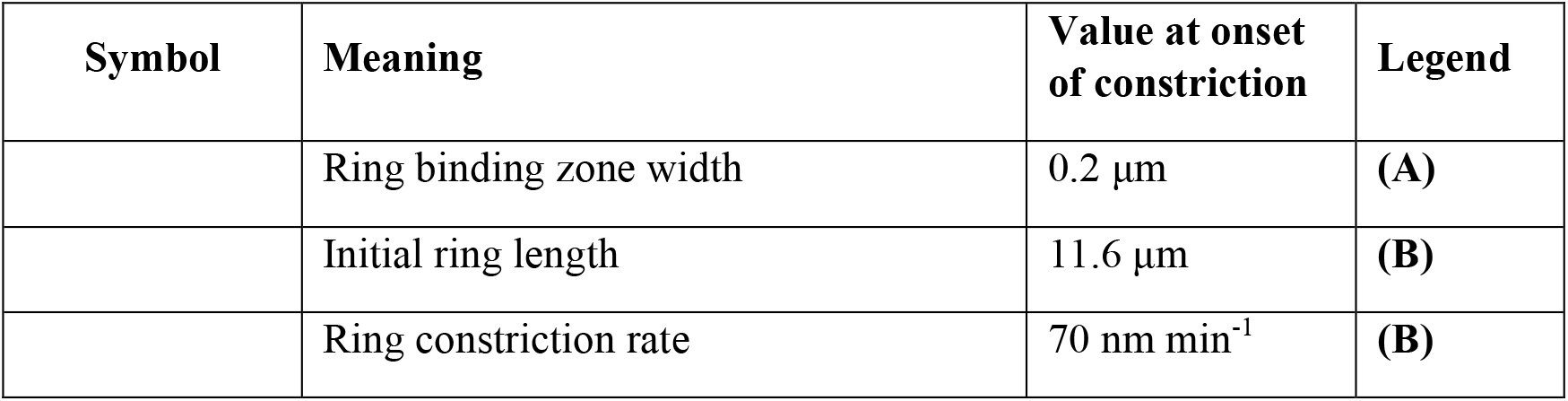
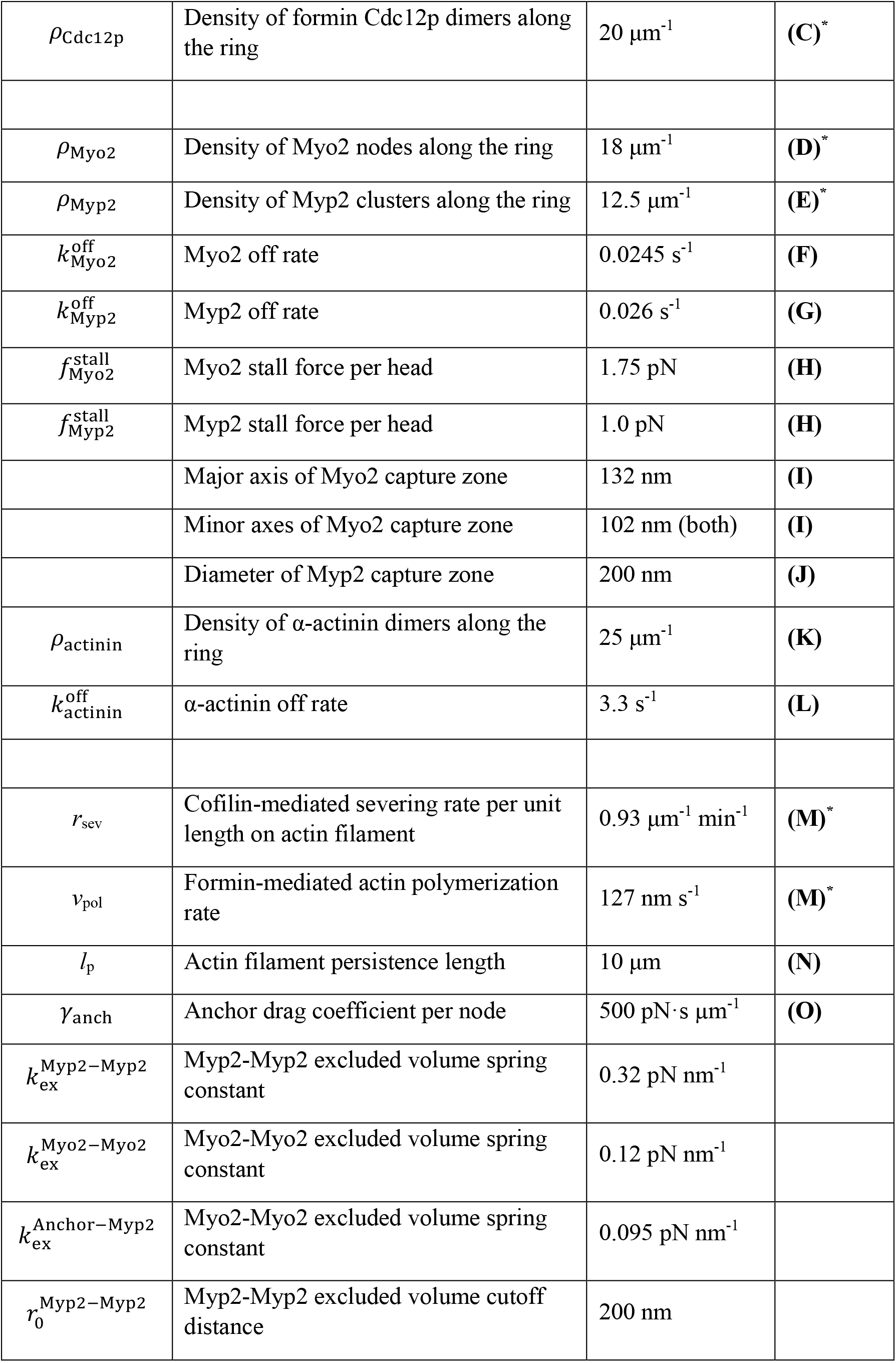
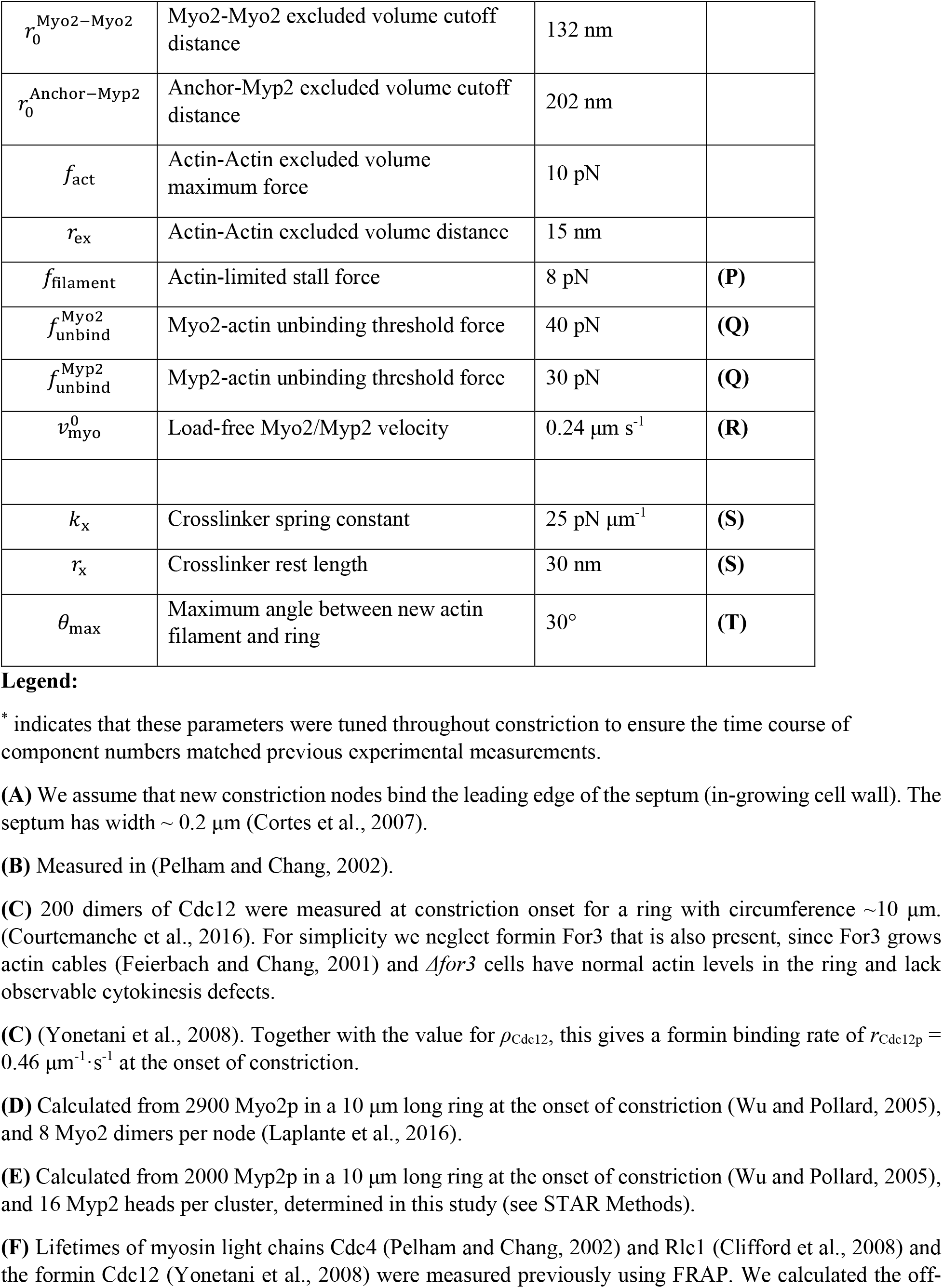
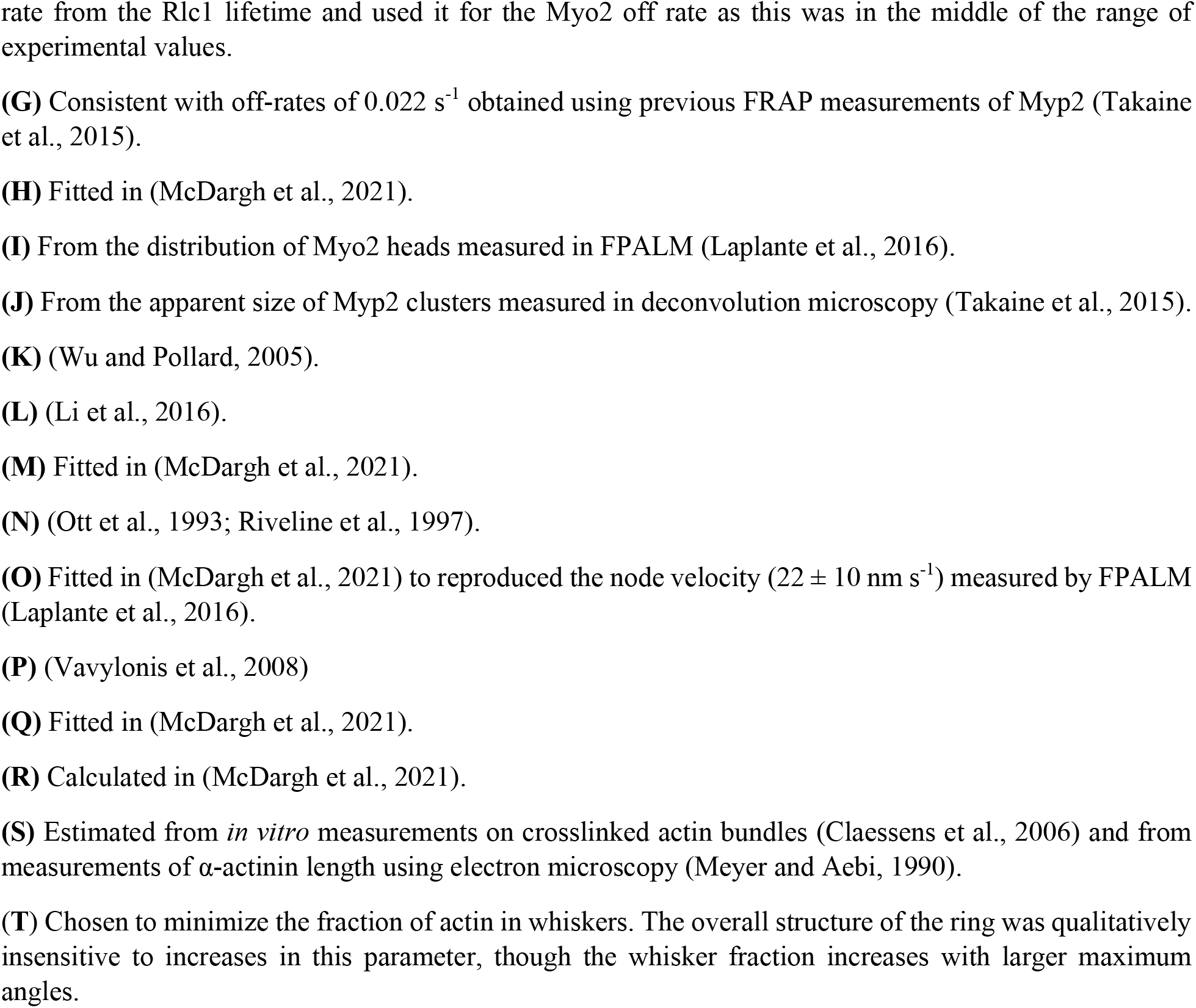
Key parameter values of the ring simulation.

### Simulations reproduce forces and structural properties of experimental contractile rings

We ran simulations of the model with a septum growing inwards at 2.4 nm s^−1^ (Thiyagarajan et al., 2015) for ∼ 25 min. Components were initially randomly positioned in a 200 nm wide circular band representing the membrane lining the inside of the septum. Constriction was initialized after a preliminary 3 min run to allow the ring to self-organize. This dynamic organization maintained itself throughout constriction and reproduced many experimental features (Fig. 2A). Consistent with experiment, the actin filaments formed a ∼ 130 nm thick bundle of ∼ 47 filaments associated with a 100 nm thick Myo2 ring overlapping a concentric 200 nm thick inner Myp2 ring, separated by ∼ 26 nm (Laplante et al., 2015; Laplante et al., 2016; McDonald et al., 2017; Swulius et al., 2018).

**Figure 2.**
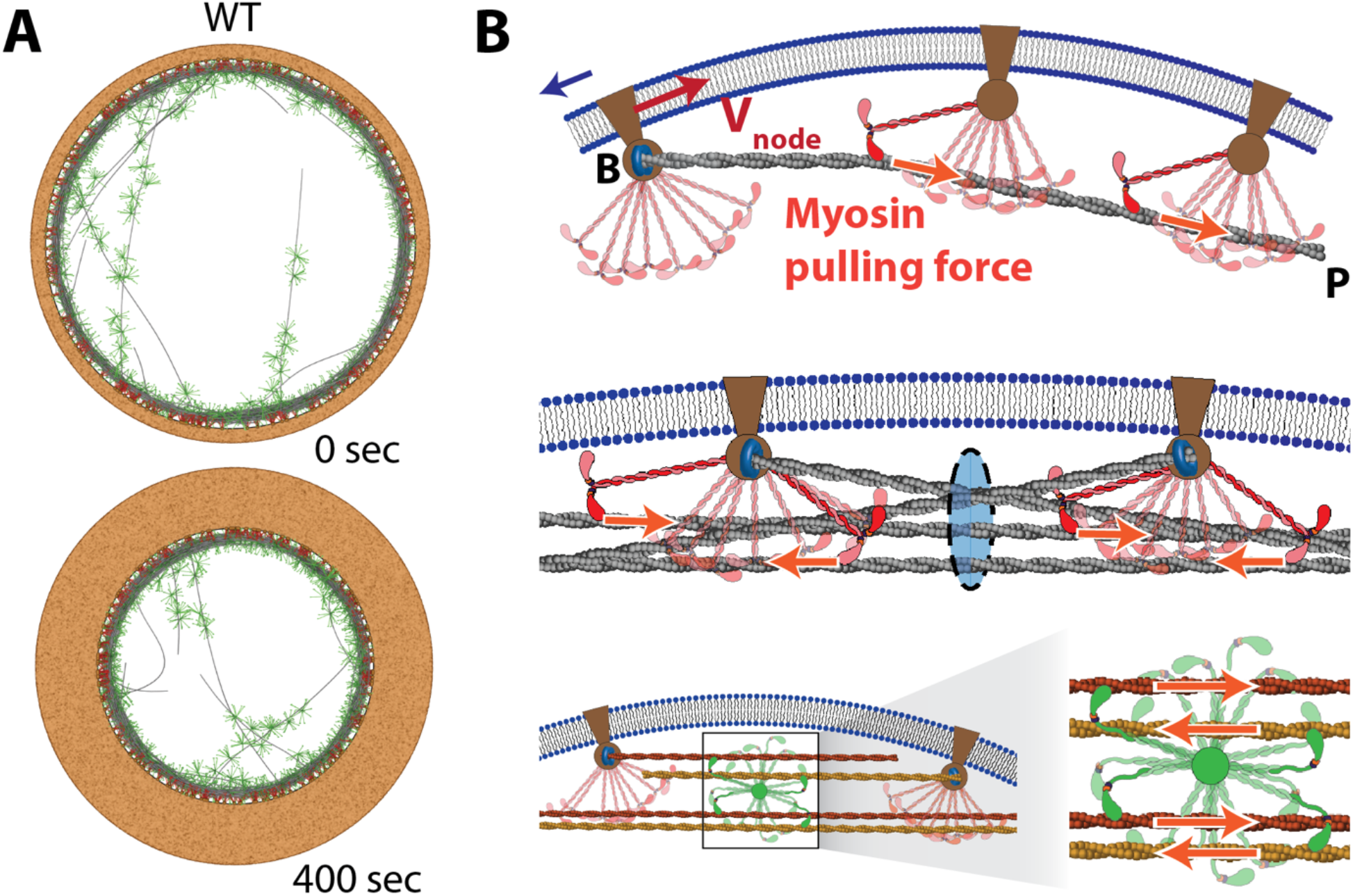
Sliding node mechanism of ring tension production. (A) Simulated wild-type contractile ring at two times during constriction. The ingrowing septum is shown as the brown region outside the ring. (B) Sliding node ring tension mechanism. Top: Individual actin filaments hosted by a node are made tense by pulling forces from Myo2 belonging to other nodes. Filament tension requires drag forces (blue arrow) that oppose motion of the node anchor relative to the membrane. Middle: the sum of the tensions of all actin filaments passing through a given cross-section of the ring gives the net ring tension. Bottom: unanchored Myp2 clusters also contribute to ring tension, by pulling on roughly equal numbers of oppositely oriented actin filaments.

Ring tension was generated by a sliding node mechanism (Fig. 2B), very different to the classic sarcomeric mechanism of striated muscle (Huxley and Niedergerke, 1954; Huxley and Hanson, 1954). When incoming nodes bound the membrane, their formins nucleated actin filaments which were pulled by Myo2 belonging to other nodes. Since the membrane-anchored nodes host ∼ 1 actin filament on average, they were pulled clockwise or counterclockwise around the ring depending on the actin orientation, with a node velocity distribution closely reproducing that measured using FPALM (Laplante et al., 2016; McDargh et al., 2021). In this way, filaments with barbed ends anchored to the membrane became tense, and a net ring tension was generated by the ∼ 200 nodes (at constriction onset) (Fig. 2B). Unanchored Myp2 clusters also contributed by pulling filaments. Ring tensions increased from ∼ 500 to ∼ 1000 pN as rings constricted (Fig. 4C), with a mean of ∼ 740 pN, close to the values measured in live fission yeast protoplasts (McDargh et al., 2021).

### Bridging instabilities are the principal structural threat to the contractile ring

The contractile ring builds a specialized organization from actin, myosin and other components that efficiently exerts centripetal force to drive constriction. For functionality, this organization must be stable over the ∼ 25 mins of constriction. Our simulations and experiments show that the biggest structural threat originates in the high tensions in filaments that must bend to form the circular bundle at the core of the ring (McDargh et al., 2021). Energetically, bent tensile filaments prefer to straighten by pulling away from the membrane into straight bridges (Fig. 3A).

**Figure 3.**
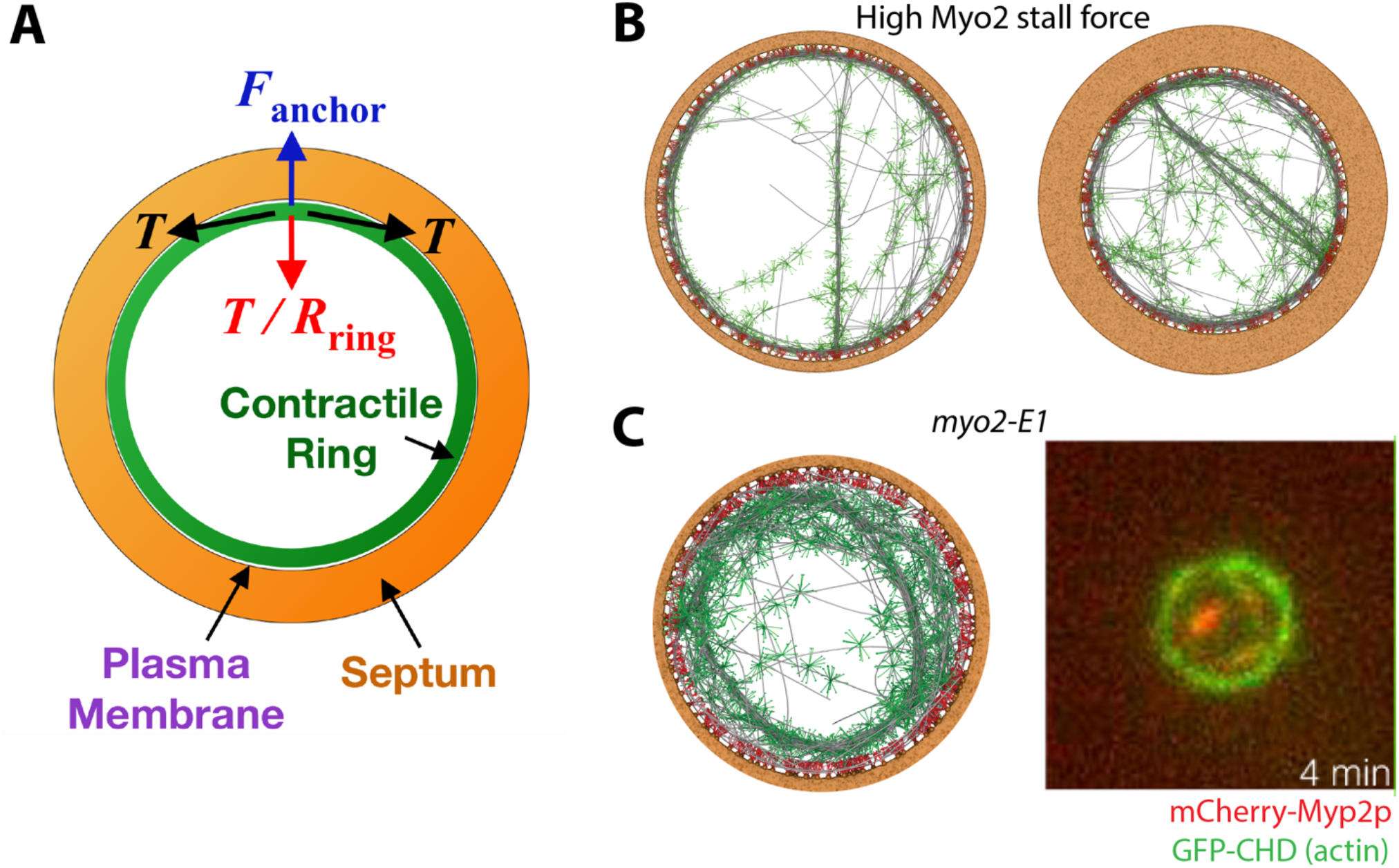
Mechanism of bridging instability. (A) The ring tension *T* produces a centripetal force per unit length *T*/*R*_ring_ for a ring of radius *R*_ring_. Normally, membrane-anchored Myo2 belonging to nodes provides the anchoring force *F*_ancher_ that opposes this force and prevents actin filaments from peeling off the membrane into straight bridges. However bridges will form if Myo2-actin binding is weak (reduced *F*_ancher_) or if the tension becomes dangerously high (increased *T*/*R*_ring_). The first scenario is realized in *myo2-E1* mutants, the second in cofilin *adf1-M3* mutants. (B) Simulations with 4-fold increased Myo2 stall force 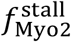 show significant bridging. (C) Simulated (McDargh et al., 2021) and experimental (Laplante et al., 2015) *myo2-E1* rings where Myo2 heads bind actin weakly show significant bridging.

Now the centripetal force per length tending to straighten a segment of a ring of radius *R*_ring_ and tension *T* into a bridge is *T*/*R*_ring_ (Fig. 3A). Further, the principal ring component anchoring actin to the membrane is the membrane-anchored Myo2, which binds actin filaments abundantly and supports them against radial displacement (McDargh et al., 2021). Myp2 makes a similar bundling contribution.

Thus we anticipate bridging will be precipitated by either (i) weakened actin-Myo2 binding, or (ii) increased ring tension *T* (higher centripetal force). Accordingly we expect bridging in simulations (i) with the temperature-sensitive Myo2-E1 mutant, which binds actin weakly and has minimal ATPase activity (Lord and Pollard, 2004; Stark et al., 2013), and (ii) with artificially elevated ring tensions *T* by using unphysiologically higher values of 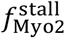 (4-fold increase), the forces exerted by Myo2 clusters (see Material and Methods). In both cases, rings suffered significant bridging, with Myp2-containing actin bridges 0.5 − 1.5 μm in length (Fig. 3B, C). The bridging in simulated *myo2-E1* rings (McDargh et al., 2021) closely matched bridging observed in live cells of the *myo2-E1* mutant, a vivid demonstration of the instability (Laplante et al., 2015) (Fig. 3C).

### Contractile rings in *adf1-M3* mutants with reduced actin severing have longer actin filaments and higher ring tension

ADF/cofilin family proteins promote actin turnover by severing actin filaments (Andrianantoandro and Pollard, 2006; Elam et al., 2013; Pavlov et al., 2007). In *adf1-M2* and *adf1-M3* mutants, the severing activity is reduced but constriction rates are normal (Cheffings et al., 2019; Chen and Pollard, 2011). To study the effects of cofilin we ran simulations to mimic the *adf1-M3* mutant. Adf1-M3 has two mutations (E132A, K133A), binds actin filaments ∼ 10-fold more slowly than wild-type and severs actin defectively (Chen and Pollard, 2011). In a severing assay (Chen and Pollard, 2011), Adf1-M3 required a ∼ 10-fold higher concentration than wild-type Adf1 for maximal actin severing activity, and even then the severing rate was ∼ 3-fold reduced from wild-type. Thus to simulate *adf1-M3* rings we set the severing rate to zero, *r*_sev_ = 0. Actin then dissociates only as whole filaments when formins dissociate from nodes (McDargh et al., 2021) (Fig. 1C).

In *adf1-M3* simulations (Fig. 4A) actin filaments were ∼ 2-fold longer than in wild-type simulations (Fig. 4B). Further, ring tension was ∼ 2-fold increased over much of the course of constriction (Fig. 4C). Given reduced severing, longer actin filaments are expected, but the enhanced tension is less obvious. To understand this, we measured tensions in individual actin filaments and found that the maximal tension at the anchored barbed end increased roughly linearly with filament length in wild-type simulations (Fig. 4D). Thus, longer filaments have higher tension, because they are bound by more myosin-II dimers. Since the total ring tension is a sum of all filament contributions over the bundle cross-section (Fig. 2B), the net ring tension is higher with longer filaments (McDargh et al., 2021).

**Figure 4.**
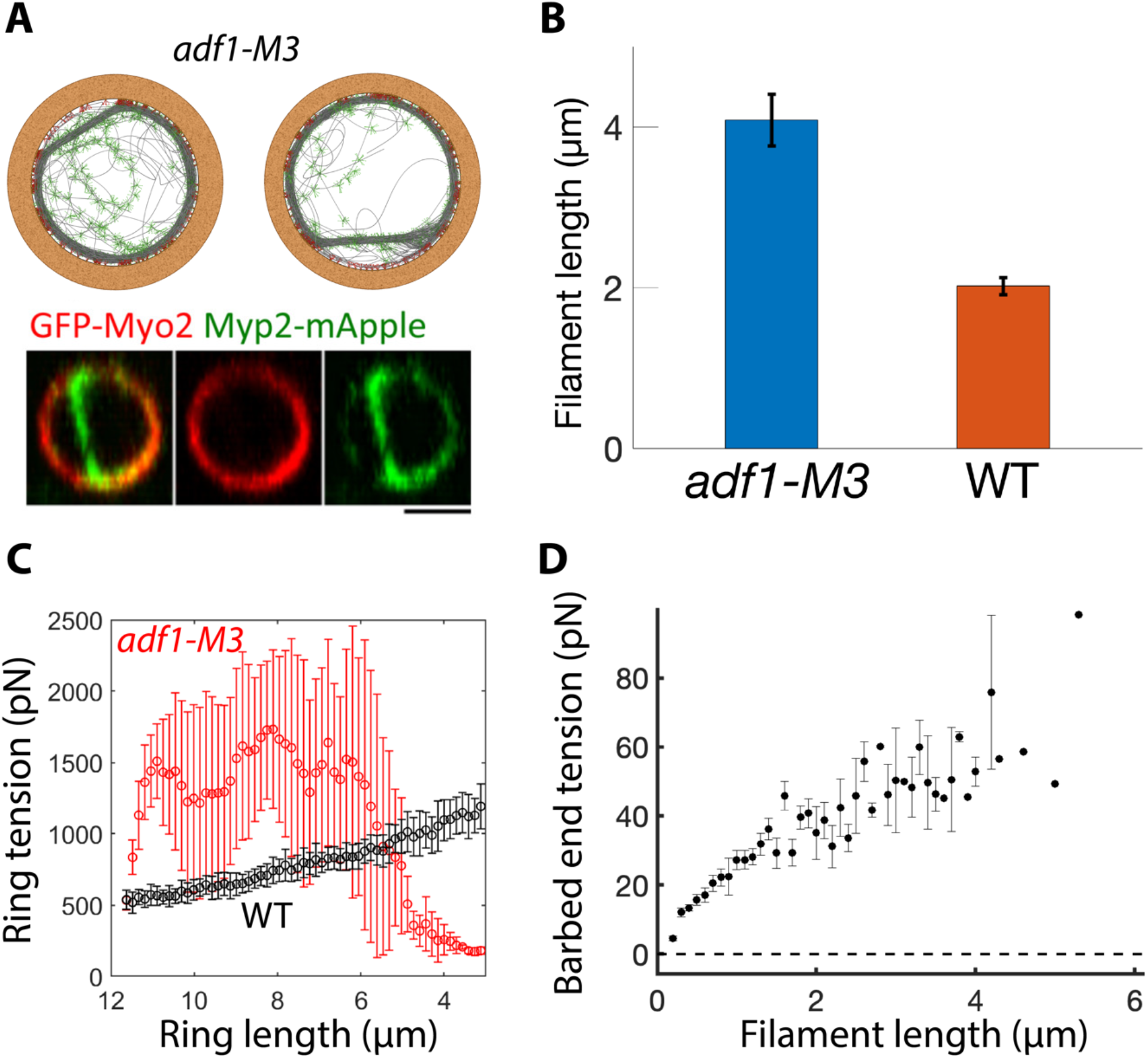
Bridging instabilities are triggered in cofilin *adf1-M3* mutants because longer actin filaments produce higher ring tension. (A) Simulated *adf1-M3* rings reproduce experimentally observed bridging (Cheffings et al., 2019). Simulated and experimental bridges contain Myp2 clusters. Scale bar: 2μm. (B) Actin filament length (mean ± SD) in simulated *adf1-M3* (*n* = 30) and wild-type (*n* = 60) rings. (C) Ring tension (mean ± SD) during constriction in simulated *adf1-M3* (*n* = 30) and wild-type (*n* = 60) rings. (D) Actin filament tension (mean ± SD) at the filament barbed end versus. filament length in simulated wild-type rings (*n* = 60).

### Simulations reproduce bridging observed in contractile rings of *adf1-M3* mutants

Interestingly, bridging instabilities are seen for the *adf1-M2* and *adf1-M3* mutants (Cheffings et al., 2019) with reduced severing activity (Fig. 4A). Why reduced severing activity triggers bridging is unknown. Our simulation results above have already provided a strong clue: longer actin filaments lead to higher ring tension *T* (Fig. 4B-D), and higher tension generates higher centripetal Laplace forces ∼ *T*/*R*_ring_ which promote bridging (Fig. 3A).

This line of reasoning was indeed confirmed by simulations, which showed dramatic bridging instabilities in cofilin mutants. The simulated bridges were remarkably similar to those observed in live cells (Fig. 4A). Leaving Myo2 behind, bridges pulled away from the membrane and comprised several actin filaments bundled by Myp2, as expected since Myp2 is unanchored and its localization to the ring depends on actin (Laplante et al., 2015; Takaine et al., 2015). Each bridge had a few tenuous attachments to the ring at the bridge ends, mediated by barbed-end-anchored actin filaments.

### Bridge formation is negatively correlated with Myo2 concentration

Consistent with experiment (Cheffings et al., 2019), bridging was more frequent in regions of lower Myo2 density around the ring. Since Myo2 is hosted by nodes, this was reflected by a negative correlation between node density and bridge location: the node density was ∼ 2-fold lower in regions of the ring where a bridge occurred than in regions without a bridge (*n* = 30 rings, Fig. 5A).

**Figure 5.**
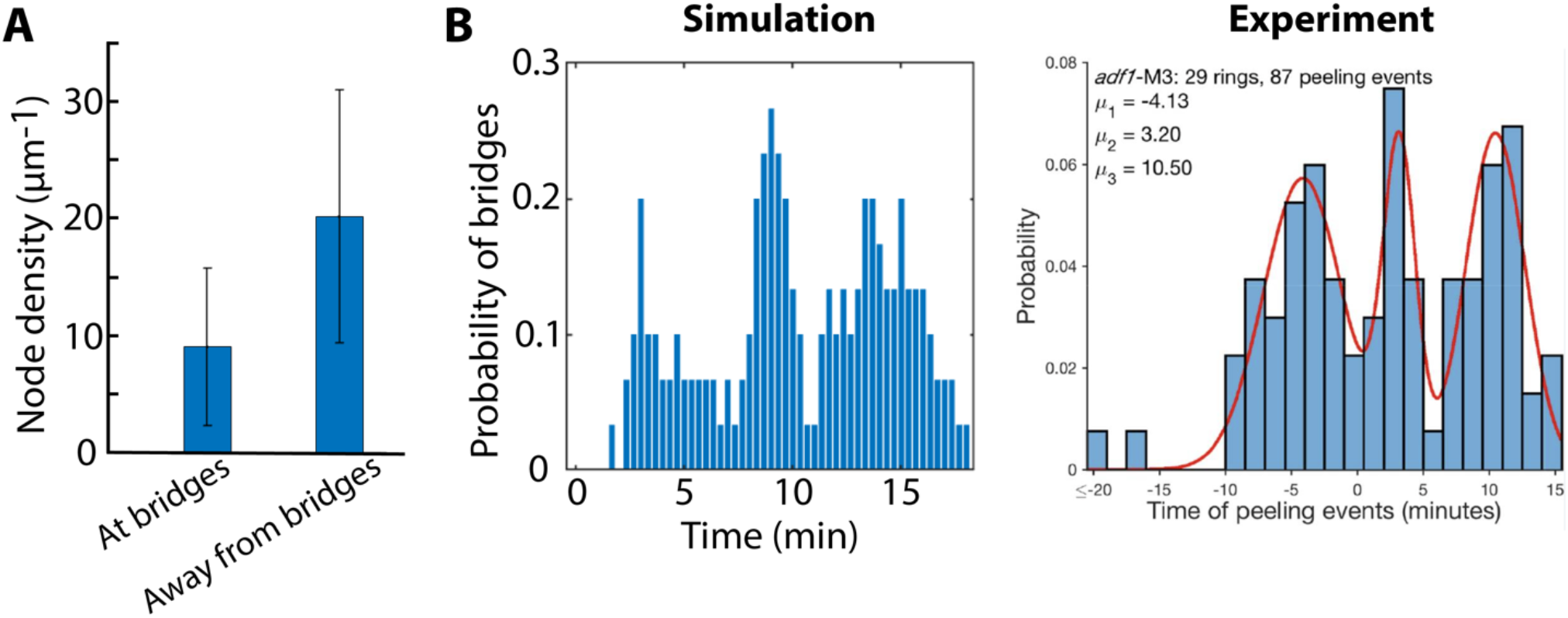
Simulated *adf1-M3* rings show negative correlation of bridging with local Myo2 concentration and cyclic bridging. (A) Node density (mean ± SD) at and away from bridging sites in simulated *adf1-M3* rings. (B) Probability of bridges at different times during constriction in simulated *adf1-M3* rings (*n* = 30) and in experimental *adf1-M3* rings (Cheffings et al., 2019).

Interestingly, this same correlation is seen in experimental images of *adf1-M3* contractile rings, where the Myo2 density is clearly lower in regions of the membrane from which a bridge has pulled away (Fig. 4A).

These observations in live cells and in simulations are consistent with bridging being a failure of Myo2-mediated anchoring. In this mechanism, tense actin filaments pull away from Myo2 that normally binds them to the membrane. Bridges thus peel away from the plasma membrane at locations where the inward Laplace forces exceed the local anchoring force from Myo2, which depends on the Myo2-actin unbinding threshold and the number of Myo2 heads that bind filaments. Thus, bridging is most likely in regions with low Myo2 density.

### Simulations reproduce cyclic bridging observed in contractile rings of cofilin mutants

In *adf1-M3* cells, bridging in contractile rings was cyclic with a period of ∼ 6 min and about 3 cycles over the course of ring constriction (Fig. 5B). Remarkably, our simulations quantitatively reproduced this behavior with considerable accuracy: ∼ 3 cycles of bridging and self-assembly occurred during constriction, each lasting ∼ 6 min (Fig. 5B).

We argue that the reproduction of this very specific experimental phenotype strongly suggests the simulation recapitulates the essential mechanisms governing the fission yeast contractile ring. In particular, we conclude that cofilin-mediated turnover that maintains actin filament lengths within a certain range are essential to protect the ring from bridging instability.

## Discussion

Here we presented evidence that actin turnover is essential for structural stability of the actomyosin contractile ring. By regulating the length of actin filaments, ADF/cofilin-mediated severing protects the ring from bridging, its principal structural threat manifested in live cells in *myo2-E1* (Laplante et al., 2015) and *adf1* mutants (Cheffings et al., 2019; Chen and Pollard, 2011). Our modeling results explain why ring disruption due to this instability is expected in these two apparently unrelated contexts: in one case due to weakened anchoring, in the other due to increased unanchoring force.

The principal job of the contractile ring is to generate tension and exert inward centripetal force that directs constriction, accomplished by myosin II pulling on the bundled curved actin filaments in the ring and making them tense. However, a curved actin filament under tension has an enormous energetic preference to become straight. A typical filament has a length *l*_act_ of the same order as its curvature, 1/*R*_ring_, where *R*_ring_∼ 1 μm is the radius of the ring. Thus its end-to-end distance increases by Δ*L*∼*R*_ring_ on straightening, giving an energy advantage −Δ*F*∼*T*_fil_ Δ*L*∼ 4000 *kT* where *T*_fil_∼ 15 pN is a typical filament tension (McDargh et al., 2021). Summed over a bundle this yields an enormous thermodynamic driving force for actin filament bundles in the ring to straighten into bridges. Thus the contractile ring is vulnerable to straightening instability and must be stabilized by anchoring. We find the principal anchoring agent is the membrane-anchored Myo2 which binds actin filaments (McDargh et al., 2021) (see Fig. 3A, C).

The contractile ring has presumably evolved to efficiently generate tension *T*, given the available myosin II in the ring. However, above a certain level tension is structurally dangerous, as the force ∼ *T*/*R*_ring_ that powers bridging will exceed the opposing Myo2-mediated anchoring forces. Cofilin-mediated severing plays a critical role because the tension depends on the actin filament length, *T*∼*ρ*_myo_ *l*_act_ (McDargh et al., 2021). Thus, for a given density *ρ*_myo_ of myosin II along the ring, longer filaments lead to higher ring tension and above a certain length provoke bridging. It is cofilin’s job to prune the actin filaments sufficiently that their length never exceeds this dangerous threshold (Fig. 4B).

Actin turnover as a necessary stabilizing process to protect the structure of actomyosin contractile assemblies is an emerging theme. Component turnover was proposed to allow the fission yeast ring to rapidly reassemble itself without trauma as it gradually constricts (McDargh et al., 2021), and actin turnover is required for homeostasis of the ring (Chew et al., 2017). In *Drosophila* embryos, actomyosin assemblies driving apical constriction were observed to recoil from adherens junctions when cofilin was inhibited by depleting cofilin phosphatase (Jodoin et al., 2015). Following breakage, the mechanical connections between the actomyosin structures and the adherens junctions were re-established on a ∼ 2 min timescale (Jodoin et al., 2015). This rupture-repair cycle is reminiscent of the ∼ 6 min bridging-repair cycles in the fission yeast contractile ring (Cheffings et al., 2019).

## Material and Methods

We used our well-tested molecularly explicit model of the contractile ring (McDargh et al., 2021). Parameters used are shown in Table 1. For simulations with artificially higher Myo2 stall force (Fig. 3B), we increased the Myo2 stall force 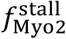 by 4 folds, from 1.75 pN to 7 pN. For simulations of *myo2-E1* mutants (Fig. 3C), we reduced the Myo2 stall force 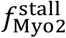 to zero, and lower the unbinding threshold 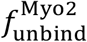 from 40 pN to 12 pN (McDargh et al., 2021). For simulations of *adf1-M3* mutants, we reduced the severing rate *r*_sev_ to zero. For other details of our molecularly explicit model, please see Ref (McDargh et al., 2021).

## Acknowledgements

We thank Mohan Balasubramanian, University of Warwick, for providing experimental images and data of *adf1-M3* rings for comparison. We thank Thomas D. Pollard, Yale University, for providing experimental images of *myo2-E1* rings for comparison. We acknowledge computing resources from Columbia University’s Shared Research Computing Facility project.

## Author Contributions

B.OS. conceived the study; Z.M. and B.OS. designed the model; Z.M., T.Z. and H.Z. performed simulations and analysis; B.OS. wrote the paper.

## Data Availability

All codes (MATLAB) of our molecular explicit model are available on our public GitHub page (https://github.com/OShaughnessyGroup-Columbia-University/constriction_of_the_fission_yeast_contractile_ring).

